# The impact of rule changes on the number and severity of injuries in the NFL

**DOI:** 10.1101/503227

**Authors:** Suril B. Sheth, Dharun Anandayuvaraj, Saumil S. Patel

## Abstract

Injuries in the National Football League (NFL) are a concern for reasons of performance, revenue, and player health. In particular, recent discoveries highlighting the long-term repercussions of concussions has drawn popular attention to the violence of the sport of American football and shone a bright light on the NFL’s efforts to reduce injuries and their long-term impact. While studies have largely focused on the impact of injuries on players’ long-term health, there has been less independent work on the evolution of injuries, namely in how the number and extent of injuries have changed over the years and what role, if any, the NFL has played in improving these figures by changing the rules or by improving protective gear technology. Here, we studied injuries from eight seasons 2010-2017, i.e. and address the impact of major rule changes that were enacted in the NFL in 2013 and 2016. We classified injuries into three major categories: arm, leg, and head, i.e. concussions. From publicly available weekly injury reports provided by each team in the NFL, we calculated the number of players who were seriously injured enough to miss at least one game of the season, and the severity of the injuries sustained, as evaluated by the number of games of the season the injured player had to sit out. The 2013 rule changes did not significantly reduce the number of players who suffered a head injury and had to sit out at least one game, and had no impact on leg or arm injuries either. Following the 2016 rule changes, we observed no change in the number of players who got injured but a significant increase in the severity of all three injury types in the two full seasons that followed. Overall, the NFL’s efforts to reduce injuries and their severity have had little success, and our findings suggest instead that rule changes or technological changes cannot curb the violence and danger present in the game today.

## Introduction

American football has come under criticism recently, however it is still Americans’ favorite sport to watch (Norman, 2018). Football has markedly slipped in terms of its popularity as compared to nearly a decade ago (Norman, 2018) and one reason could be the negative publicity football has received regarding injuries in general and concussions in particular (Mez et al., 2017) and the long-term debilitating effect injuries are likely to have on player health and quality of life (there might be other reasons as well). As a result, a bright spotlight has been placed in recent years on the National Football League (NFL) – the premier professional association of American football – and the steps the NFL has taken and is taking to protect players, reduce injuries, and improve the long-term impacts of these injuries on player health.

Two ways that the NFL can achieve player safety are to institute new rules or new technology. Rule changes have many underlying causes, e.g. more points, better television (TV) ratings, more dollars, or better public relations, i.e. PR. One of the purported causes is player safety. Technological changes such as changes in protective helmets, pads etc. are made with a view to, in large part, improving player safety. It is imperative that new metrics be developed that will examine the efficacy of these rules and/or changes in technology in terms of limiting player injuries and enhancing player safety.

Major injuries that occur to players in the NFL are the following – knee and ankle injuries, e.g. a torn anterior cruciate ligament (ACL) or medial cruciate ligament (MCL), shoulder and arm injuries, e.g. a dislocated shoulder, and concussions, i.e. an injury to the brain caused by a single or repeated hits to the head and neck area leading to the brain being shaken in the skull. In brief, major injuries are injuries to the arm, leg, and head.

Injuries negatively impact the short-term health of NFL players as well as their long-term health and playing career. Let us take the example of concussions. Casson et al. (2003) analyzed the videos of all concussions between the 1996 and 2001 NFL seasons, and concluded that concussions are caused by hits to the head that occur at rapid speeds (roughly 9.3 m/s on average). NFL athletes may not feel the effects of concussions sustained in their playing career immediately, but their effects are likely to be detrimental to their health in the long term.

According to Didehbani, Munro Cullum, Mansinghani, Conover, & Hart (2013), who compared 42 retired NFL players with a control group of non-NFL players, players who have sustained concussions during their playing career will be more likely to have depression as they become older: scores in the Beck Depression Inventory (BDI-II) (Beck, Ward, Mendelson, Mock, & J., 1961; Steer, Ball, Ranieri, & Beck, 1997) in all three categories – cognitive, affective, and somatic – were higher in NFL players than in the control cohort. As compared to former NFL players who have not had a concussion in their playing career, former NFL players with three or more concussions in their playing career are five times more likely to have mild cognitive impairment in the future and three times more likely to have memory problems in the future (Guskiewicz et al., 2005). Guskiewicz et al. (2005) found that recurrent cerebral concussions in former NFL players were associated with an earlier onset stage for Alzheimer’s disease, and Omalu et al. (Omalu et al., 2006; Omalu et al., 2005) found that recurrent cerebral concussions in former NFL players were associated with other brain issues such as MCI and Parkinson’s disease. Leg injuries are major injuries as well and can have impact on the long-term as well as the short-term. Myer et al. (2011) showed that, as compared to athletes who have no injury history of a torn ACL, the athleticism and speed of athletes who have returned to compete in their sport before one year following a torn ACL is worse. Erickson et al. (2014) showed that the possibility of a quarterback returning to play another game in the NFL after a torn ACL injury is extremely high but that there is a small decline in the quality of their play.

Not only do injuries affect the players’ health and performance, injuries also hinder the ability of the NFL to expand its market to other countries, such as Germany, Mexico, and England (Vrentas, 2015). Generally speaking, healthy teams, i.e. teams that have fewer injuries to its players than the league average, make the playoffs and ones that are unhealthy do not, e.g. 7 of the 10 healthiest NFL teams in the 2008 NFL season made the playoffs as compared to only 2 of the 10 unhealthiest NFL teams (Barnwell, 2009). Tom Brady, reigning league MVP and star quarterback for the New England Patriots, a team that had made it all the way to the Super Bowl the previous season, suffered a season-ending torn ACL, and the New England Patriots did not make the playoffs that season (Barnwell, 2009). Thus, injuries impact team success and as well as the bottom line, and shedding light on what the NFL is doing to reduce injuries as well as their impact is of critical importance.

The NFL makes rule changes every year to make the game more exciting, increase TV viewership and live audiences, and enhance player safety and reduce injuries. In keeping, rule changes made for the 2013 and 2016 seasons particularly stand out as in those years, crucial rule changes were made with an eye towards enhancing player safety and preventing injury. There were several important safety related rule changes made before the 2013 season (NFLFootballOperations, 2013). First, players were prohibited from initiating contact with the crown of their helmet outside the tackle box, the idea being that the rule change would reduce concussions. Second, rule violations regarding contact to the head of defenseless players would now be subject to suspension, which should reduce further the occurrence of concussions. Third, peel back blocks which were previously illegal outside the tackle box were now considered illegal anywhere on the field, a change which was made to reduce knee injuries to defensive players. Finally, all players besides kickers and punters were now required to wear knee and thigh pads, which would presumably reduce the number of leg injuries and/or their severity. In addition, the NFL announced the introduction of a set of concussion assessment guidelines in 2013, which were developed by the league’s Head, Neck and Spine Committee in an attempt to better detect concussions during practices and games (Flynn, 2016; NFL’sHeadNeckandSpineCommittee, 2013). There were two important safety related rule changes made before the 2016 season (NFLFootballOperations, 2016). First, all chop blocks were made illegal, which was a popular change for defensive players, who long experienced knee and ankle injuries in those situations. Second, horse-collar tackle rules were expanded to include the area at the name plate or above when grabbed to pull a runner toward the ground – the play was made illegal if the runner was pulled to the ground or his knees were buckled by the action because the opposing player grabbed the inside collar or the jersey at the name plate. Both rule changes were made to reduce ankle and knee injuries. On the basis of these health and safety related rule changes, we expect sharp declines in the number of players who suffered injuries - head and leg injuries, in particular – in the seasons following the 2013 and 2016 rule changes as compared to the seasons prior.

The aforementioned studies have analyzed the impact of various injuries on player performance, overall short-term and long-term health, the player’s team’s success, and even the league’s bottom line but do not mention whether the number and severity of these changes has changed over the years, nor do they discuss the effectiveness of preventative measures such as rule changes on the number and extent of said injuries. The NFL has published a study that gives the total number of concussions and ACL and MCL injuries in years 2012-2015, but no details have been provided, such as how many players were injured, whether or not they had to sit out any games because of the injuries, and, if they were out of action, for how many weeks (NFL.com, 2016). Although the NFL has taken actions in reducing the quantity and severity of these injuries by enforcing rule changes and adoption of new technology to protect players, one still questions how impactful these rule changes have been in protecting NFL players.

Our study is among the first to examine this very issue using data that are publicly available: do rule changes have an impact on injury? In particular, we will examine whether rule changes instituted by the NFL in 2013 and in 2016 reduced injuries to the arm, leg, and head and/or diminished their severity.

## Materials & Methods

We collected data on the number of major injuries in the NFL from the past eight full NFL regular seasons: 2010-2017 (see “http://www.fftoday.com/nfl/10_injury_wk1.html,” for an example. There are 17 such injury reports for each NFL season.). We did not include the postseason in the data because only 11 out of 32 NFL teams participate in the playoffs, and the sample set would be uneven. We did not include the preseason in the data because teams use the preseason as a warm up to the regular season; therefore, NFL players, especially starters, do not play as hard in the preseason as in the regular season. In short, for our first set of analyses, we counted the number of players across the league who suffered an injury that caused them to miss at least one game of the regular season. Note that irrespective of how many weeks a given player remains out because of a given injury, he counts as one toward the first measure.

For a second, complementary set of analyses, we evaluated the severity of each major injury, and our measure was the number of weeks a player was on the injury report, representing the time a player was sidelined (note that the NFL does not use existing technology to measure the impact of hits; our measure of evaluating the number of games missed is a reasonable way of evaluating severity of injury). For a given season, we obtained the weekly injury report of each of the 32 NFL teams, followed the injury history of each player, and thus obtained a measure for the severity of his injury by calculating the number of weeks that the player was out of action. We then obtained a league-wide average across all 32 NFL teams for a given season. Thus, for each season, we had two measures – the number of players across the league who had to sit out at least one game (week), and the number of weeks the player had to sit out because of the injury. In addition, we focused on three major classes of injury – leg, arm, and head – in order to see if and how rule changes differentially impacted injuries to different body parts. Leg injuries included injuries to the knee, calf, ankle, hamstring, thigh, and fibula, arm injuries included injuries to the shoulder, forearm, and elbow, and head injuries included injuries to the head and eye.

As mentioned in the Introduction above, the rule changes instituted by the NFL in 2013 and 2016 were benchmarks. For each of the two measures described above, we statistically compared – using two-tailed unpaired *t*-tests – the following: seasons 2010-2012 versus seasons 2013-2015, and seasons 2010-2015 versus seasons 2016-2017, in order to address the impact of the 2013 and 2016 rule changes respectively.

## Results

Figs. 1A and 1B respectively show the number of players whose leg injuries were severe enough that the injured player had to miss at least one regular season game, and the mean number of games players missed for seasons 2010-2017 – a measure of the severity of their leg injury. To address the impact of the 2013 rule changes, we compared a) the number of injured players (*t*(4) = − 0.402, *p* = 0.708, not significant, or ns) and b) mean number of games missed (*t*(4) = −0.600, *p* = 0.581, ns) in seasons 2010-2012 before the rule change versus seasons 2013-2015 after. As Fig. 1A and 1B show, and statistics confirm, the 2013 rule changes did not have a statistically distinguishable impact on leg injuries. A similar analysis was done to address the impact of the 2016 rule changes. Comparing seasons 2010-2015 before the rule change and seasons 2016-2017 after, there was no significant change in the number of players who suffered leg injuries after the rule change (*t*(6) = 0.776, *p* = 0.467, ns), but there was a significant *increase* in the number of weeks the player had to miss game time because of said injury (*t*(6) = −23.61, *p* = 0.000000379). Thus, the 2016 rule changes did not have a positive impact on leg injuries, in terms of reducing their number or severity.

**Fig. 1.**
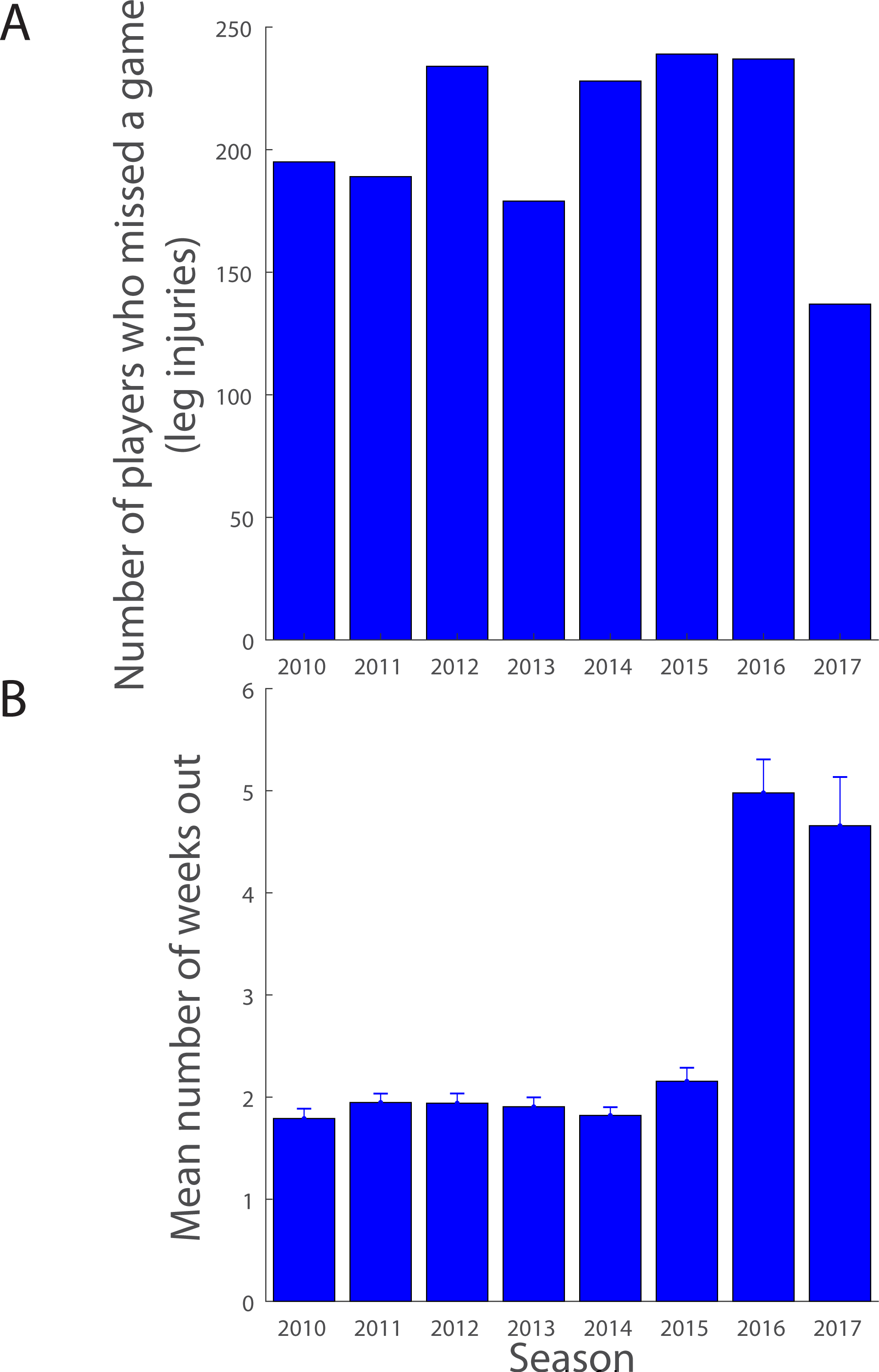
Number and severity of leg injuries (A) shows a bar graph of the total number of players who missed at least one regular season game because of a leg injury (ordinate) as a function of NFL season (abscissa; 2010-2017). (B) shows a bar graph of the severity of the leg injury in terms of the mean number of weeks, or games of the regular season, the injured player was unable to play (ordinate) as a function of NFL season (abscissa; 2010-2017). Error bars are one standard error of the mean (s.e.m).

To address the impact of the 2013 rule changes on arm injuries, we compared seasons 2010-2012 versus seasons 2013-2015 for a) the number of players injured (*t*(4) = −1.051, *p* = 0.353, ns) and b) mean number of games missed because of the injury (*t*(4) = −0.587, *p* = 0.589, ns). As Fig. 2A and B show, and statistics confirm, the 2013 rule changes did not positively impact arm injuries, statistically or numerically. We did a similar analysis of the 2016 rule changes. Comparing seasons 2010-2015 before the rule change and seasons 2016-2017 after, there was a statistically insignificant change in the number of players who suffered arm injuries after the rule change (*t*(6) = −0.887, *p* = 0.409, ns), but a highly significant *increase* in the number of weeks the player had to miss game time because of said injury (*t*(6) = −13.287, *p* = 0.0000112). Thus, the 2016 NFL rule changes did not reduce the number or severity of arm injuries to NFL players.

**Fig. 2.**
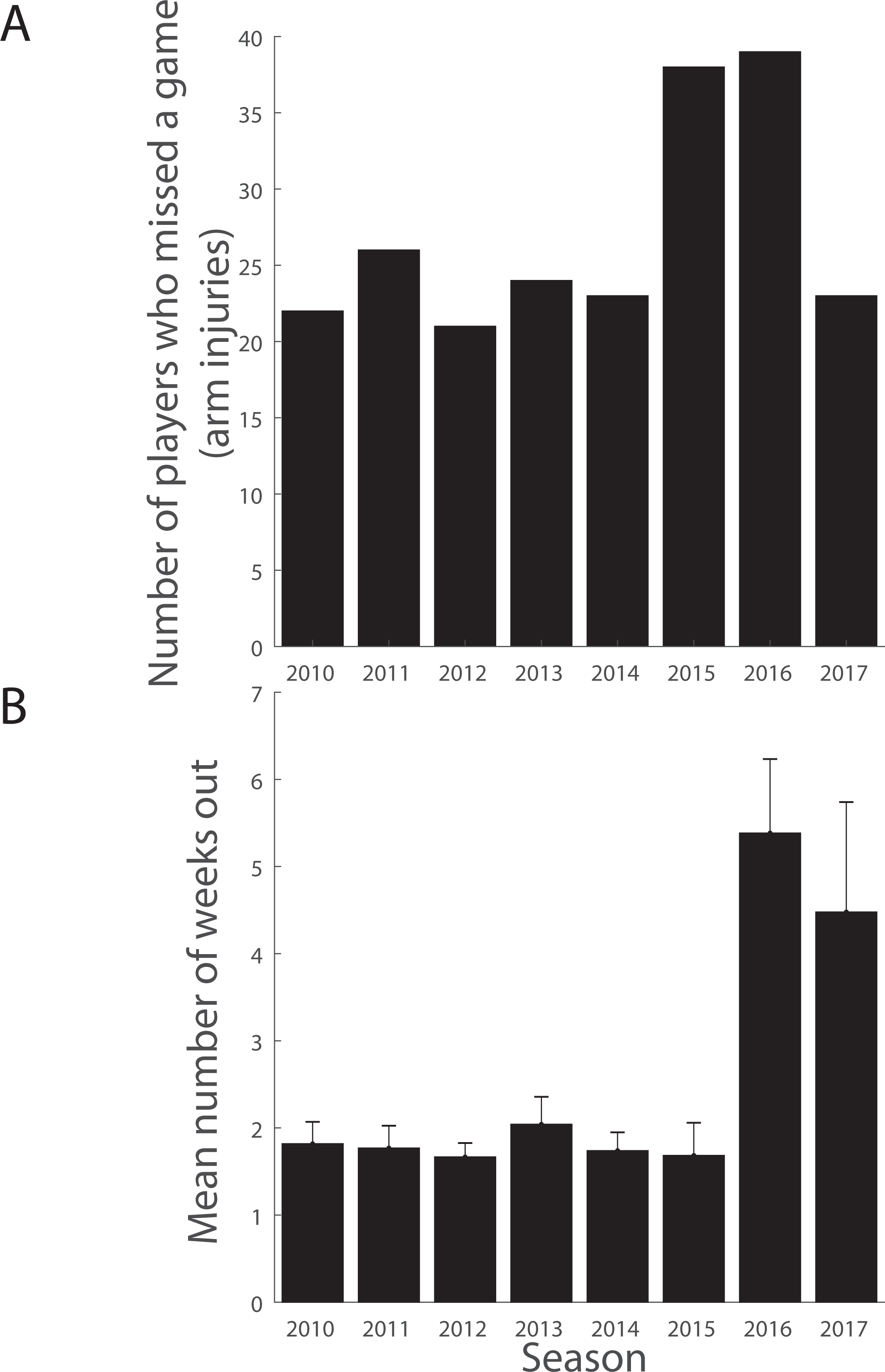
Number and severity of arm injuries (A) shows a bar graph of the total number of players who missed at least one regular season game because of an arm injury (ordinate) as a function of NFL season (abscissa; 2010-2017). (B) shows a bar graph of the severity of the arm injury in terms of the mean number of weeks, or games of the regular season, the injured player was unable to play (ordinate) as a function of NFL season (abscissa; 2010-2017). Error bars are one standard error of the mean (s.e.m).

To address the impact of the 2013 rule changes on head injuries (concussions), we compared seasons 2010-2012 versus seasons 2013-2015 for a) the number of injured players (*t*(4) = 0.753, *p* = −0.338, ns), and b) mean number of games missed (*t*(4) = 0.052, *p* = 0.961, ns). As Fig. 3A and B show, and statistics confirm, the 2013 rule changes did not statistically reduce the number of head injuries (concussions) or reduce the severity, at least in terms of the number of weeks the injured player lost. We did a similar analysis of the 2016 rule changes. Comparing seasons 2010-2015 before the rule change and seasons 2016-2017 after, there was no statistically significant change in the number of players who suffered head injuries following the 2016 rule change (*t*(6) = −0.998, *p* = 0.357, ns) but a significant *increase* in the number of weeks the player had to miss game time because of them (*t*(6) = −11.102, *p* = 0.0000318).

**Fig. 3.**
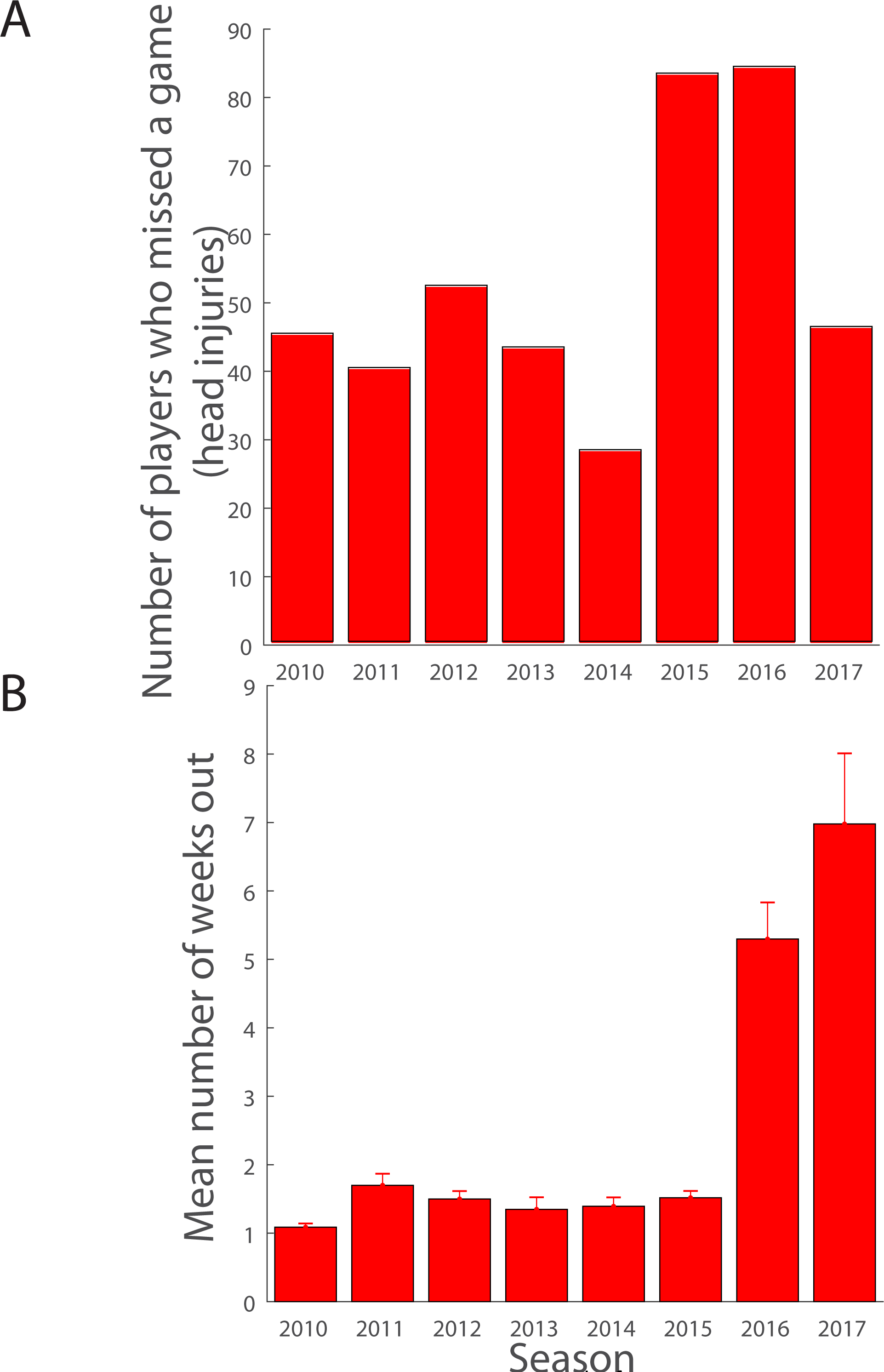
Number and severity of head injuries (A) shows a bar graph of the total number of players who missed at least one regular season game because of a head injury (ordinate) as a function of NFL season (abscissa; 2010-2017). (B) shows a bar graph of the severity of the head injury in terms of the mean number of weeks, or games of the regular season, the injured player was unable to play (ordinate) as a function of NFL season (abscissa; 2010-2017). Error bars are one standard error of the mean (s.e.m).

In summary, the 2013 or the 2016 rule changes had no positive impact on leg, arm, or head injuries. Neither the 2013 rule changes nor the 2016 rule changes reduced the number of players who suffered concussions. All the results, including ones showing increase in the severity of leg, arm and head injuries following the 2016 rule changes, are discussed below along with possible explanations and implications.

## Discussion

Our research aimed to study whether the NFL’s efforts – through rule changes and educational efforts – to reduce player injuries has had a positive impact. In particular, we focused on the major rule changes implemented in the NFL just before the 2013 and 2016 seasons, and we studied the evolution in the number and severity of three major classes of injuries – leg, arm, and head (concussions) through seasons 2010-2017. Our findings are significant to the sport of American football as important life and death questions have arisen regarding the importance of player safety to the NFL and the quality of the precautions that the NFL has taken in recent years. Our study is among the first to look at this issue independent of the NFL.

Our study showed that neither the 2013 nor the 2016 rule changes had a positive impact on injuries to leg, arm, or head. The 2013 rule changes did not reduce the number of players who suffered injuries to the leg, arm, or head; neither did the 2016 rule changes. In particular, the rule changes made in 2013 targeted concussion safety, and yet the number of players who suffered head injuries (concussions) did not go down significantly, and the severity of those players who did suffer a head injury – as measured by the number of games they were out – remained unaffected by the rule changes. Furthermore, compared to seasons past, the 2016 and 2017 seasons showed a sharp, highly significant uptick in the severity of arm, leg, and head injuries, as measured by the average number of games (weeks) the injured player was unavailable to play. Thus, there is an overall increase in the severity of all injuries in the last two full NFL seasons. We discuss the findings, possible underlying causes, and implications below.

We begin with a negative finding, i.e. there is not a steadily improving record in reducing the number of players suffering leg and arm injuries over the years, or, once an injury occurs, in reducing its severity. Our results suggest that the series of well-intentioned changes, namely increased emphasis on reducing player-to-player contact, and tinkering of the rules to purportedly enhance player safety, has not had a positive impact on arm and leg injuries. One argument has been that shoulder pads have not improved much over the years while the game has become more violent with stronger, better players. In the inaugural season of the NFL in 1920, shoulder pads were large, thick, heavy, and noticeable despite being covered by a jersey (Longman, 2014); today, shoulder pads are smaller, roughly 50% lighter, and could barely be noticed through a jersey (Longman, 2014). The new shoulder pads as well as the absence of knee/thigh pads may has made the modern player faster, more agile, and better at blocking and tackling (Longman, 2014), but the tradeoff may well be an increase in the number and/or severity of shoulder and arm injuries. Our findings support the skeptics – the severity of arm (includes shoulder) injuries has increased dramatically and significantly over the last two full NFL seasons.

Concussions are a slightly different story. The NFL made a number of changes prior to the 2013 season targeted towards enhancing concussion safety: instituted rules such as banning helmet-to-helmet hits, making kickoff plays safer, and limiting the amount of contact allowed in practices. Hand in hand, it adopted new guidelines for sideline evaluation, rules on preseason education, baseline testing and the establishment of personnel – specifically an NFL physician advisor unaffiliated with any NFL team – to conduct concussion evaluations. However, these rule changes failed in significantly reducing the number of players with concussions who missed a game (Fig. 3A). One counter-argument is that the new concussion guidelines, which call for a minimum of 29 medical professionals in every game, increased the probability of detecting a concussion, so that the actual number of concussions went down but the probability of detecting one when it occurred went up – thus, the two probabilities counteract each other. This is a real possibility, albeit an unlikely one. Indeed, if the probability of detecting a concussion increased, low-grade concussions will also be detected (i.e. a lax criterion level in a signal detection theory paradigm implies a lower threshold); therefore the average severity of the concussion will be lower – Fig. 3B shows and statistics confirm this was not the case.

In this context, the sharp and significant increase in concussion severity in the last two full NFL seasons (2016, 2017; Fig. 3B) runs counter to the NFL’s efforts and needs explaining. One possible reason for the recent uptick in concussions is that with recent news about concussions and their long-term impact, the NFL is now exercising a high degree of caution before placing their concussed players back on the field. The alternative, arguably likelier, reason is that concussions are more severe than before: NFL players today are faster, bigger, and stronger than before (NFLFootballOperations, 2018), which could possibly contribute to the more severe head injuries, and consequently longer durations of player inactivity (as an aside, note that helmet to helmet contacts are only one cause of concussions, helmet to body and helmet to ground contacts are common causes as well, and no new rules have been adopted to minimize helmet to ground contact). In keeping, the increased momentum resulting from these changes in body structure (and/or the enhanced muscle mass of the modern NFL player may be at the upper limit of what the tendons and ligaments can bear, which results in more fragile arms and legs, and increased likelihood for injuries to be more severe) should lead to more severe injuries *of all kinds*: our data showing a significant uptick in leg (Fig. 1B) and arm (Fig. 2B) injuries over the same period as the significant uptick in head injuries (Fig. 3B) are in agreement with this reasoning. According to others who have looked at NFL injuries, more serious knee injuries every year means more serious head injuries (Breslow, 2013). We contend there is a second, additional contributing factor to the uptick: the gradual evolution of the game from a defensive, running game to a more exciting offense-dominant, passing oriented game over the past decades, which inevitably means more plays per game (compare e.g. SportingCharts, 2009; 2016; significantly more plays in 2016 NFL season vs. 2009 NFL season (p < 0.001); see site for additional evidence), more players involved in each play (due to more movement at the line of scrimmage as well as downfield), and more acrobatic plays. It is reasonable to expect that the above two factors – evolutions of player and game – combined have had a negative impact on player safety, and more severe leg, arm, and head injuries are an inevitable consequence.

The NFL’s own assessment is that their concussion protocols are working based on data that they began sharing with the public since 2012 (NFL.com, 2016). A lower number of diagnosed concussions is a sign of progress, but so is a higher number as per the NFL, because the NFL claims that a higher number points to a sign of “culture change”, a point that was made in 2016 (Campbell, 2016) and again, back in 2010 when the number of concussions first showed an uptick compared to the year before (NFL.com, 2010; Wagner, 2016). Such flexible interpretations of data muddy the use of the number of concussions, and injuries in general, as a measure of progress in enhancing player safety. The numbers provided by the NFL include a large majority of concussions that occur during the game (or practice) but do not lead to any loss of playing time in future games. That is to say, a concussed player was back on the field for their team’s very next game. One reason for the quick turnaround (except for the major concussions which do require a prolonged period of recovery and downtime for the player) is that the NFL and its team of traumatic brain injury physicians, with its new awareness of concussions, has become better at concussion management and speeding up recovery. Although we are not aware of scientific breakthroughs to help people recover from concussions, the possibility is a real one, nonetheless. An alternative possibility is based on a different interpretation of concussion management. The concussion protocol calls for the concussed player to be examined and monitored in the training room on a daily basis. The periodic nature of the tests, conducted by the team neuropsychology consultant (note: *not* an independent consultant), could lead to the player learning to game the protocol and hasten an apparent recovery. The “learning to the test” approach (we do not know if there was any “teaching to the test” as well) has been found to exist in a number of disciplines including education (Strathern, 1997), economics (Goodhart, 1981), and academic research (Biagioli, 2016) and is a real possibility here as well. At present, we are unable to distinguish between the two possibilities and further work is required.

Our study is among the first to objectively report on and analyze injuries in the NFL using sources other than the NFL, but it has limitations. The websites used to gather data on the number and severity of major injuries do not list the injury reports for teams who have a bye week – a week where the team does not play a game – which leads to an undercount of the actual number of injuries. This also underestimates the measure of injury severity, because a player can get injured the week before the bye week and the bye week spent recovering from the injury does not count. Additionally, all injuries do not get reported by the NFL teams. Moreover, injuries in which the player did not have to spend any time out of the game were not counted in our study. Finally, injuries that occur late in a season may have smaller severity values because injuries after the regular season is over are not monitored. Nevertheless, these limitations, while important and lead to an under-reporting of injuries in general, do not systematically bias the results of our study.

## Conclusions

We studied the effect of rule changes made by the NFL before the 2013 and 2016 seasons. The 2013 rule changes failed to reduce the number of players with concussions leading to a loss of game time, the number of players who suffered arm and leg injuries leading to a loss of game time, or the severity of leg, arm, or head injuries as evaluated by the number of weeks the player had to miss. Following the 2016 rule changes, yet again, there was no reduction in the number of players who had leg, arm, or head injuries; however, we observed a significant increase in the severity of leg, arm, and head injuries. Despite the NFL’s tinkering with the rules and the continuous evolution of pads and protective gear technology over the years to purportedly enhance player safety, we have not observed a decreasing trend over the years in terms of the number of players who got injured or in the number of weeks the injured player has to recover–in fact, the opposite. Our findings support the idea that the NFL game is inherently too violent, and are consistent with the assertion that rule changes or technological changes in protective gear are unable to meaningfully reduce the number and severity of injuries in the NFL. This may be a sobering conclusion for NFL players’ health and the economic health of the NFL going forward. From a bird’s-eye view, our work highlights the importance of data over theory, technology, and marketing in sports research.

## Acknowledgments

The research was conducted as part of an AP Research class at Carnegie Vanguard High School (SBS). We thank the AP Research teacher, Ms. Hill for her constant encouragement and help on the project.

## References

Barnwell, B. (2009, 9.12.2009). Some N.F.L. Injuries May Hurt More Than Others, The New York Times. Retrieved from https://www.nytimes.com/2009/09/13/sports/football/13injuries.html

Beck, A. T., Ward, C. H., Mendelson, M., Mock, J., & J., E. (1961). An inventory for measuring depression. Arch. Gen. Psychiatry, 4(6), 561–571.

Biagioli, M. (2016). Watch out for cheats in citation game. Nature, 535(7611), 201. doi: 10.1038/535201a

Breslow, J. (2013). For the NFL, Focus on Concussions Yields Mixed Results. Frontline. https://www.pbs.org/wgbh/frontline/article/about-concussion-watch-2/

Campbell, R. (2016). Reported concussions in NFL up 32 percent in 2015. The Chicago Tribune. https://www.chicagotribune.com/sports/football/ct-nfl-concussions-spt-0130-20160129-story.html

Didehbani, N., Munro Cullum, C., Mansinghani, S., Conover, H., & Hart, J., Jr. (2013). Depressive symptoms and concussions in aging retired NFL players. Arch Clin Neuropsychol, 28(5), 418–424. doi: 10.1093/arclin/act028

Erickson, B. J., Harris, J. D., Heninger, J. R., Frank, R., Bush-Joseph, C. A., Verma, N. N.,… Bach, B. R. (2014). Performance and return-to-sport after ACL reconstruction in NFL quarterbacks. Orthopedics, 37(8), e728–734. doi: 10.3928/01477447-20140728-59

Flynn, E. (2016). What is the NFL’s concussion protocol? Sports Illustrated. from https://www.si.com/nfl/nfl-concussion-protocol-policy-history

Goodhart, C. (1981). Problems of Monetary Management: The U.K. Experience. In A. S. Courakis (Ed.), Inflation, Depression, and Economic Policy in the West (pp. 111–146). Lanham, MD: Rowman & Littlefield.

Guskiewicz, K. M., Marshall, S. W., Bailes, J., McCrea, M., Cantu, R. C., Randolph, C., & Jordan, B. D. (2005). Association between recurrent concussion and late-life cognitive impairment in retired professional football players. Neurosurgery, 57(4), 719–726; discussion 719-726. http://www.fftoday.com/nfl/10_injury_wk1.html. xfrom http://www.fftoday.com/nfl/10_injury_wk1.html

Longman, J. (2014, 1.30.2014). Shoulder Pads Slim Down in Faster, Sleeker N.F.L., The New York Times. Retrieved from https://www.nytimes.com/2014/01/31/sports/football/shoulder-pads-slim-down-in-faster-sleeker-nfl.html

Mez, J., Daneshvar, D. H., Kiernan, P. T., Abdolmohammadi, B., Alvarez, V. E., Huber, B. R.,.. McKee, A. C. (2017). Clinicopathological Evaluation of Chronic Traumatic Encephalopathy in Players of American Football. JAMA, 318(4), 360–370. doi: 10.1001/jama.2017.8334

Myer, G. D., Schmitt, L. C., Brent, J. L., Ford, K. R., Barber Foss, K. D., Scherer, B. J.,… Hewett, T. E. (2011). Utilization of modified NFL combine testing to identify functional deficits in athletes following ACL reconstruction. J Orthop Sports Phys Ther, 41(6), 377–387. doi: 10.2519/jospt.2011.3547

NFL.com. (2010). Concussions reported in NFL up 21 percent from last season. from http://www.nfl.com/news/story/09000d5d81cdf2d6/article/concussions-reported-in-nfl-up-21-percent-from-last-season

NFL.com. (2016). 2015 Injury Data. from http://static.nfl.com/static/content/public/photo/2016/01/29/0ap3000000629781.pdf

NFL’sHeadNeckandSpineCommittee. (2013). NFL Head, Neck and Spine Committee’s Protocols Regarding Diagnosis and Management of Concussion. 1-6. Retrieved from http://static.nfl.com/static/content/public/photo/2013/10/01/0ap2000000254002.pdf website:

NFLFootballOperations. (2013). Health and Safety Rules Changes.

NFLFootballOperations. (2016). 2016 Rule Changes and Points of Emphasis.

NFLFootballOperations. (2018). Evolution of the NFL Player.

Norman, J. (2018). Football still Americans’ favorite sport to watch. https://news.gallup.com/poll/224864/football-americans-favorite-sport-watch.aspx

Omalu, B. I., DeKosky, S. T., Hamilton, R. L., Minster, R. L., Kamboh, M. I., Shakir, A. M., & Wecht, C. H. (2006). Chronic traumatic encephalopathy in a national football league player: part II. Neurosurgery, 59(5), 1086-1092; discussion 1092-1083. doi: 10.1227/01.NEU.0000245601.69451.27

Omalu, B. I., DeKosky, S. T., Minster, R. L., Kamboh, M. I., Hamilton, R. L., & Wecht, C. H. (2005). Chronic traumatic encephalopathy in a National Football League player. Neurosurgery, 57(1), 128-134; discussion 128-134.

SportingCharts. (2009). Team Total Offense and Defense Plays Per Game: 2009 NFL Season. from https://www.sportingcharts.com/nfl/stats/team-total-offense-and-defense-plays-per-game/2009/

SportingCharts. (2016). Team Total Offense and Defense Plays Per Game: 2016 NFL Season. from https://www.sportingcharts.com/nfl/stats/team-total-offense-and-defense-plays-per-game/2016/

Steer, R. A., Ball, R., Ranieri, W. F., & Beck, A. T. (1997). Further evidence for the construct validity of the Beck depression Inventory-II with psychiatric outpatients. Psychol Rep, 80(2), 443–446. doi: 10.2466/pr0.1997.80.2.443

Strathern, M. (1997). ‘Improving ratings’: audit in the British University system. European Review, 5, 305–321.

Vrentas, J. (2015). The NFL’s Future in Europe. Sports Illustrated. https://www.si.com/mmqb/2015/07/24/nfl-future-europe

Wagner, L. (2016). NFL Report: Concussion Diagnoses Increased 32 Percent. https://www.npr.org/sections/thetwo-way/2016/01/29/464880358/nfl-report-concussion-diagnoses-increased-32-percent

